# Propagation of a rapid cell-to-cell H_2_O_2_ signal over long distances in a monolayer of cardiomyocyte cells

**DOI:** 10.1101/2023.12.19.572374

**Authors:** Yosef Fichman, Linda Rowland, Thi Thao Nguyen, Shi-Jie Chen, Ron Mittler

## Abstract

Cell-to-cell communication plays a cardinal role in the biology of multicellular organisms. H_2_O_2_ is an important cell-to-cell signaling molecule involved in the response of mammalian cells to wounding and other stimuli. We previously identified a signaling pathway that transmits wound-induced cell-to-cell H_2_O_2_ signals within minutes over long distances, measured in centimeters, in a monolayer of cardiomyocytes. Here we report that this long-distance H_2_O_2_ signaling pathway is accompanied by enhanced accumulation of cytosolic H_2_O_2_ and altered redox state in cells along its path. We further show that it requires the production of superoxide, as well as the function of gap junctions, and that it is accompanied by changes in the abundance of hundreds of proteins in cells along its path. Our findings highlight the existence of a unique and rapid long-distance H_2_O_2_ signaling pathway that could play an important role in different inflammatory responses, wound responses/healing, cardiovascular disease, and/or other conditions.

**Highlights:**

- Wounding induces an H_2_O_2_ cell-to-cell signal in a monolayer of cardiomyocytes.
- The cell-to-cell signal requires H_2_O_2_ and O_2_·^-^ accumulation along its path.
- The signal propagates over several centimeters changing the redox state of cells.
- Changes in the abundance of hundreds of proteins accompanies the signal.
- The cell-to-cell signal requires paracrine and juxtacrine signaling.

**Graphical Abstract:** 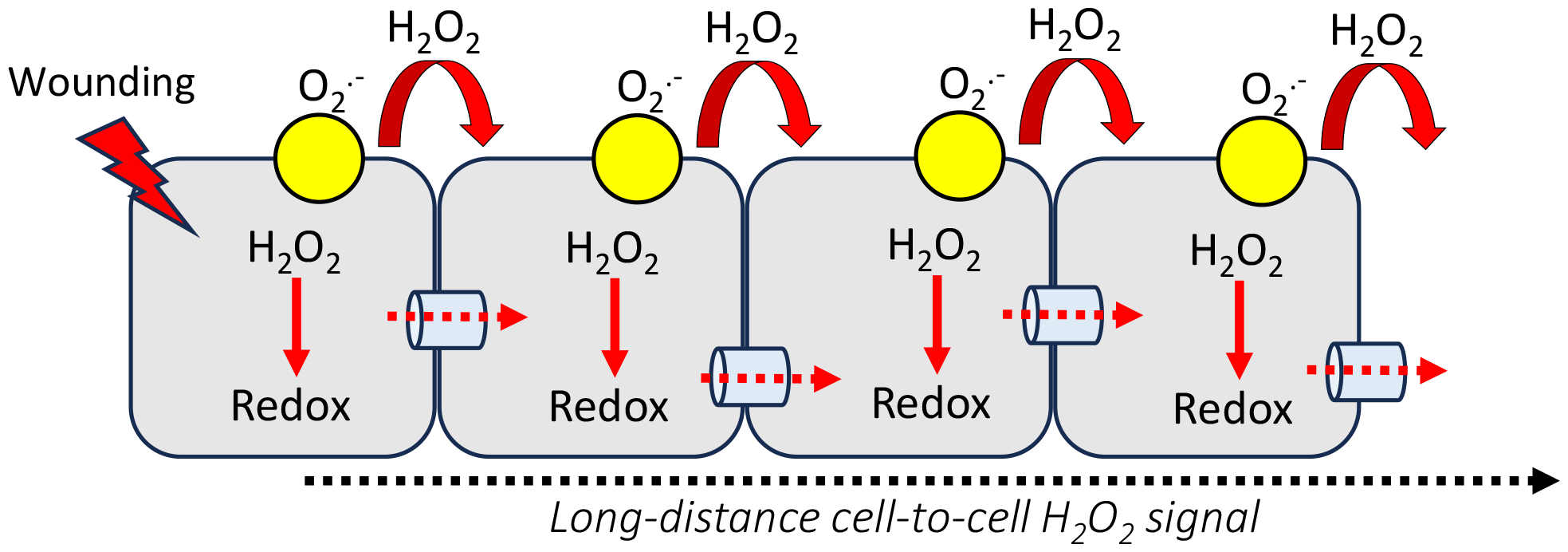

## Introduction

Cell-to-cell communication, including paracrine, juxtacrine, and autocrine signaling, plays a cardinal role in the development, maintenance, repair, and milieu/stimuli responses of multicellular organisms. Among the many signals that mediate cell-to-cell signaling, H_2_O_2_ plays a key role as a regulator of cellular and extracellular redox responses^1-4^. In most cell-to-cell H_2_O_2_ signaling studies to date, the production of H_2_O_2_ is mediated by plasma membrane localized NADPH oxidase (NOX)/dual oxidase (DUOX) in response to cellular changes in calcium levels, and the O_2_·^-^ produced extracellularly is converted to H_2_O_2_, either by superoxide dismutases (SODs), or by the peroxidase domain of DUOXs^5-11^. Extracellular H_2_O_2_ can then enter the cells that produced it, or neighboring cells, through peroxiporins and coordinate their responses to the stimuli^1-4^. H_2_O_2_ packed in exosomes, or proteins secreted by one cell, can also trigger H_2_O_2_ responses in neighboring cells^12-14^. H_2_O_2_ signals produced by different cells have multiple functions. For example, in response to wounding, H_2_O_2_ produced at the injury boundary generates a gradient that attracts leukocytes to the site of injury^15-17^. In other cases, H_2_O_2_ sensed by neighboring cells via redox-regulated receptors triggers in the receiving cells multiple responses that include changes in calcium levels and the activation kinases^18^. H_2_O_2_ can also alter the redox state of the producing and receiving cells, and/or the extracellular matrix, and trigger multiple responses^1-4^.

Overall, unless H_2_O_2_ is packed into exosomes, it is believed that the distance an extracellular H_2_O_2_ signal can travel in mammalian tissues is in the range of 100-200 μm, and potentially up to 1 mm^15-17,19^. This limitation is mainly attributed to the H_2_O_2_ scavenging activity of the extracellular matrix/neighboring cells, that generates a gradient from the producing cells, all the way up to the point H_2_O_2_ levels drop to normal levels at the tissue(s) they are generated at^15-17,19^. Recent studies in plant and mammalian cells have, however, challenged this concept showing that H_2_O_2_ signals can propagate within minutes over long distances (measured in centimeters) in different plant tissues, as well as in monolayers of human MDA-MB-231 breast cancer and rat H9c2 cardiomyocyte cells, and isolated mice hearts^18,20-25^. The mechanism that was proposed to drive this ‘long-distance’ H_2_O_2_ signal was termed the reactive oxygen species wave (‘ROS wave’) and it is based on the enhanced production of H_2_O_2_ by each cell along the path of the signal. An initiating cell would therefore produce H_2_O_2_ that is sensed by a neighboring cell and cause it to actively generate H_2_O_2_, triggering a cascade of cell-to-cell ‘enhanced H_2_O_2_ production’ state^18,20-25^. In this respect, it is important to note that it is not H_2_O_2_ that diffuses over long distances, rater, it is a state of ‘enhanced H_2_O_2_ production’ that is propagating in a cell-to-cell fashion from the initiating cell(s) through the tissue, sometimes in a directional manner, with a rate of 0.15-0.4 cm/min^18,20-25^. Although H_2_O_2_ does not diffuse over long distances *per se* in this cell-to-cell signaling mechanism, it is nonetheless required for the signal to propagate, as external application of catalase, NOX inhibitors, or the introduction of a loss-of-function NOX mutation into cells, block it^18,20-25^.

Although the ROS wave has been extensively studied in plants, only one report described it in monolayers of mammalian cells, as well as other eukaryotic organisms such as algae and amoeba (living in a community or film)^20^. Moreover, changes in protein/mRNA expression associated with the ROS wave were only determined in plants and algae^20,21,26^. Many questions regarding this mode of long-distance H_2_O_2_ signaling in mammalian cells remain, therefore, unanswered. Here, we provide evidence that the long-distance H_2_O_2_ signal in a monolayer of cardiomyocyte cells is accompanied by changes in protein abundance, enhanced cytosolic H_2_O_2_ levels, and altered cytosolic redox state, along its path, and that it depends on extracellular production of superoxide and gap junction function.

## Materials and Methods

H9c2(2-1) rat cardio myoblast (CRL-1446; ATCC) cells were plated at a density of 500,000/well for 6-well plates, and 25,000/well for 96-well plates and imaged using IVIS (Revvity) or Lionheart FX (BioTek), accordingly^20,27^. roGFP2 and roGFP2-Orp1 expressing cells were transfected with pMF1707 and pMOS016 (Addgene), accordingly, grown, and imaged as described previously^24,28-30^. Peroxy Orange 1, Dihydroethidium (DHE), CellROX orange, superoxide dismutase, catalase, APX115, and Setanaxib were obtained and used as described previously^20,30^. Carbenoxolone (CBX) and β-glycyrrhetinic acid (βGA) (Sigma) were added 60 min before imaging. Injury, imaging, and data analyses were performed as described in^20^. Protein extraction was performed from H9c2(2-1) cells grown on 100 mm plates treated and imaged with the IVIS platform as described in^20,28,29^. Cells were scrapped from the plates and proteins were extracted, identified (Uniprot ID-UP000002494), and analyzed as described in^20,28,29^. Proteomics data are available vin ProteomeXchange with identifier PXD047131. Statistical analysis was performed as described in^20^.

## Results and Discussion

### The rapid long-distance H_2_O_2_ signal is accompanied by changes in protein abundance in cells along its path

Monolayers of H9c2 Rat cardiomyocytes cells were injured and cells 0.1-1 cm away from the injury site (termed ‘local’ cells), as well as cells 3-4 cm away from it (termed ‘systemic’ cells), were sampled from injured, and uninjured (control) monolayers (Fig. 1A). In parallel, changes in H_2_O_2_ and O_2_·^-^ levels in response to the injury were measured (Fig. 1A). Proteomics analysis of cells sampled from the local and systemic locations revealed that, compared to control unwounded cells from these locations, over 3400 and 1800 proteins significantly changed in their abundance in the local and systemic cells, in response to wounding, respectively (Figs. 1B, S1; Tables S1-S3). Importantly, over 870 proteins that changed in abundance in local cells also changed in abundance in systemic cells (Fig. 1B; Table S3). These proteins could be directly associated with the ‘enhanced H_2_O_2_ production state’ triggered in the local and systemic cells in response to wounding (Fig. 1A). This group of proteins was enriched in molecular chaperone, oxidoreductase acting on sulfur, and membrane insertase activities (Fig. S1C), and included proteins such as ferredoxin, thioredoxin, MnSOD, catalase, peroxiredoxin, ferritin, and different proteins involved in glutathione metabolism (Table S3), suggesting that they are associated with altered redox/H_2_O_2_ levels in cells. An overlap between the proteins identified in response to wounding by our study and zebrafish proteins altered in response to H_2_O_2_ ^31^ was also found (Fig. 3C). The identified proteins/pathways associated with the state of enhanced H_2_O_2_ production (Figs. 1, S1; Tables S1-S3) could be used in future studies to dissect the process of long-distance cell-to-cell H_2_O_2_ signaling. Although it would be ideal to study the proteomics responses of H9c2 monolayers treated with an external antioxidant, or composed of NOX/DOUX mutants, these treatments/manipulations would affect both the initiation as well as the propagation of the H_2_O_2_ signal and would not provide a high-resolution analysis of this process. Instead, methods to generate monolayers composed of sections that contain NOX/DOUX mutations flanked by sections that do not (like our studies in *Chlamydomonas*)^20^ should be developed. In addition to imaging H_2_O_2_ levels with the IVIS apparatus (Fig. 1A), we also used an automated fluorescent microscope to measure the rates of ROS accumulation in cells along the path of the signal (Fig. S2). This analysis revealed that the highest rates of ROS accumulation occurred close to the injury site, followed by lower rates at the middle and opposite edge of the long-distance H_2_O_2_ signal path (Fig. S2). This analysis further supported the model of enhanced H_2_O_2_ production by different cells along the path of the signal^4,20,21^.

**Fig. 1.**
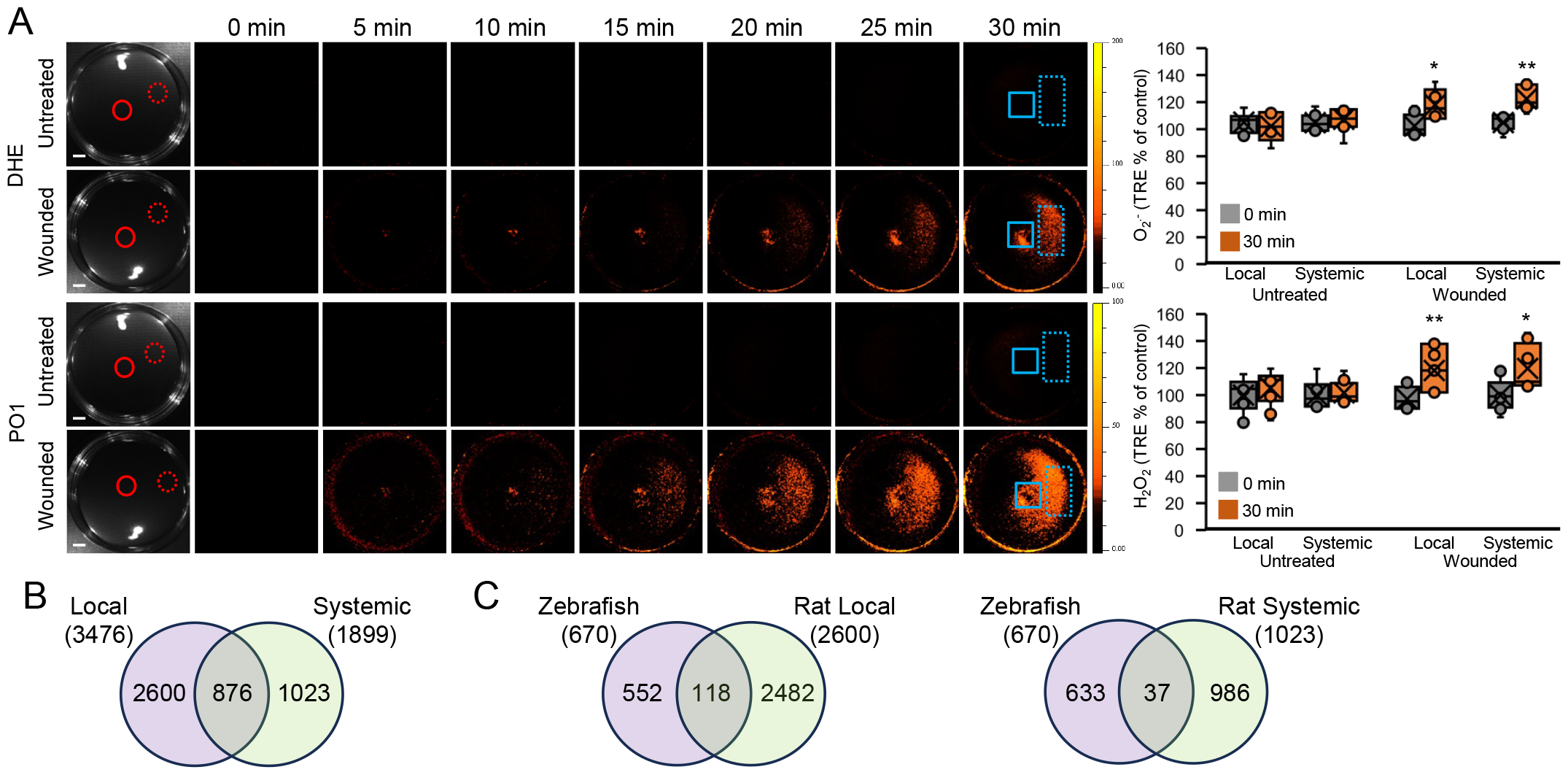
The wound induced H_2_O_2_ accumulation signal is accompanied by changes in protein abundance in cells along its path. **A**. Monolayers of rat cardiomyocytes (H9c2) grown in culture were wounded with a heated metal rod to induce injury (average injury diameter is 120 μm)^20^ and O_2_·^-^ and H_2_O_2_ accumulation were imaged using DHE and PO1 respectively. Representative time-lapse images of whole plate O_2_·^-^ and H_2_O_2_ accumulation in treated and untreated plates are shown alongside bar graphs of combined data from all plates used for the analysis at the 0- and 30-min time points (local and systemic; Background levels of O_2_·^-^ and H_2_O_2_, measured at time 0, were subtracted from all other time points). Local and systemic sampling locations are indicated with solid and dashed circles, respectively. **B**. Venn diagrams showing changes in protein abundance in local and systemic cells (Following the experimental design shown in A; Local and systemic sampling locations are indicated with solid and dashed rectangles, respectively), 30 min following wounding of the local cells as described above. **C**. Venn diagrams showing overlap between the proteins that changed in abundance in local (Left) or systemic (right) rat cardiomyocytes (B) and proteins that changed in abundance in response to H_2_O_2_ application in Zebrafish^31^. All experiments were repeated at least 3 times with 10 plates per experiment. Data is presented in A as box plot graphs; X is mean ± S.E., N=30, **P < 0.01, *P < 0.05, Student t-test. Scale bar, 2 cm. Abbreviations: DHE, dihydroethdium; PO1, peroxy orange 1; TRE, total radiant efficiency.

**Fig. 2.**
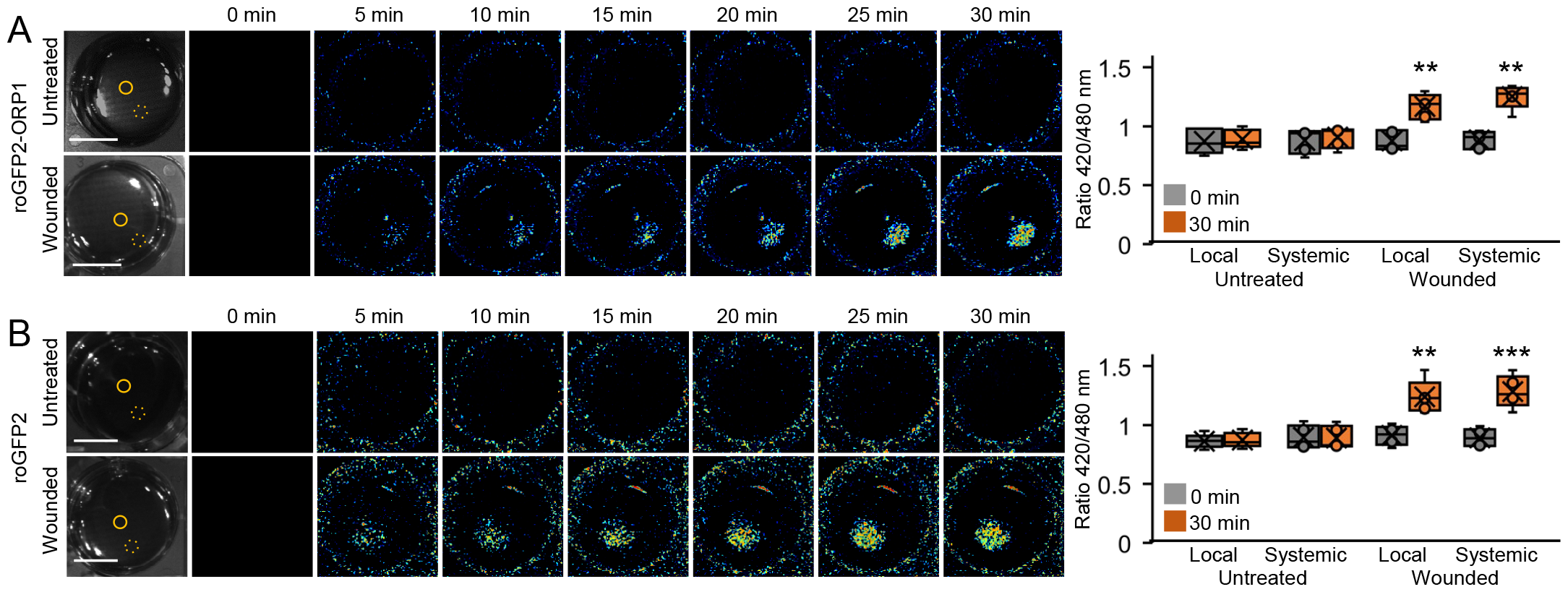
The wound induced H_2_O_2_ accumulation signal is accompanied by changes in cytosolic H_2_O_2_ and redox levels. **A**. Monolayers of rat cardiomyocytes (H9c2) expressing roGFP2-Orp1 grown in culture were wounded with a heated metal rod to induce injury^20^ and cytosolic H_2_O_2_ levels were imaged as the ratio between 420/480 nm^24^. Representative time-lapse images of whole plate changes in cytosolic H_2_O_2_ levels in treated and untreated plates are shown alongside bar graphs of combined data from all plates used for the analysis at the 0- and 30-min time points (local and systemic; Background levels of cytosolic H_2_O_2_ levels, measured at time 0, were subtracted from all other time points). **B**. Same as A, but for rat cardiomyocytes (H9c2) expressing roGFP2. Calibration of the roGFP2-Orp1 and roGFP2 signals in rat cardiomyocytes are shown in Fig. S3. All experiments were repeated at least 3 times with 10 plates per experiment. Data is presented in A and B as box plot graphs; X is mean ± S.E., N=30, **P < 0.01, ***P < 0.001, Student t-test. Scale bar, 1 cm.

**Fig. 3.**
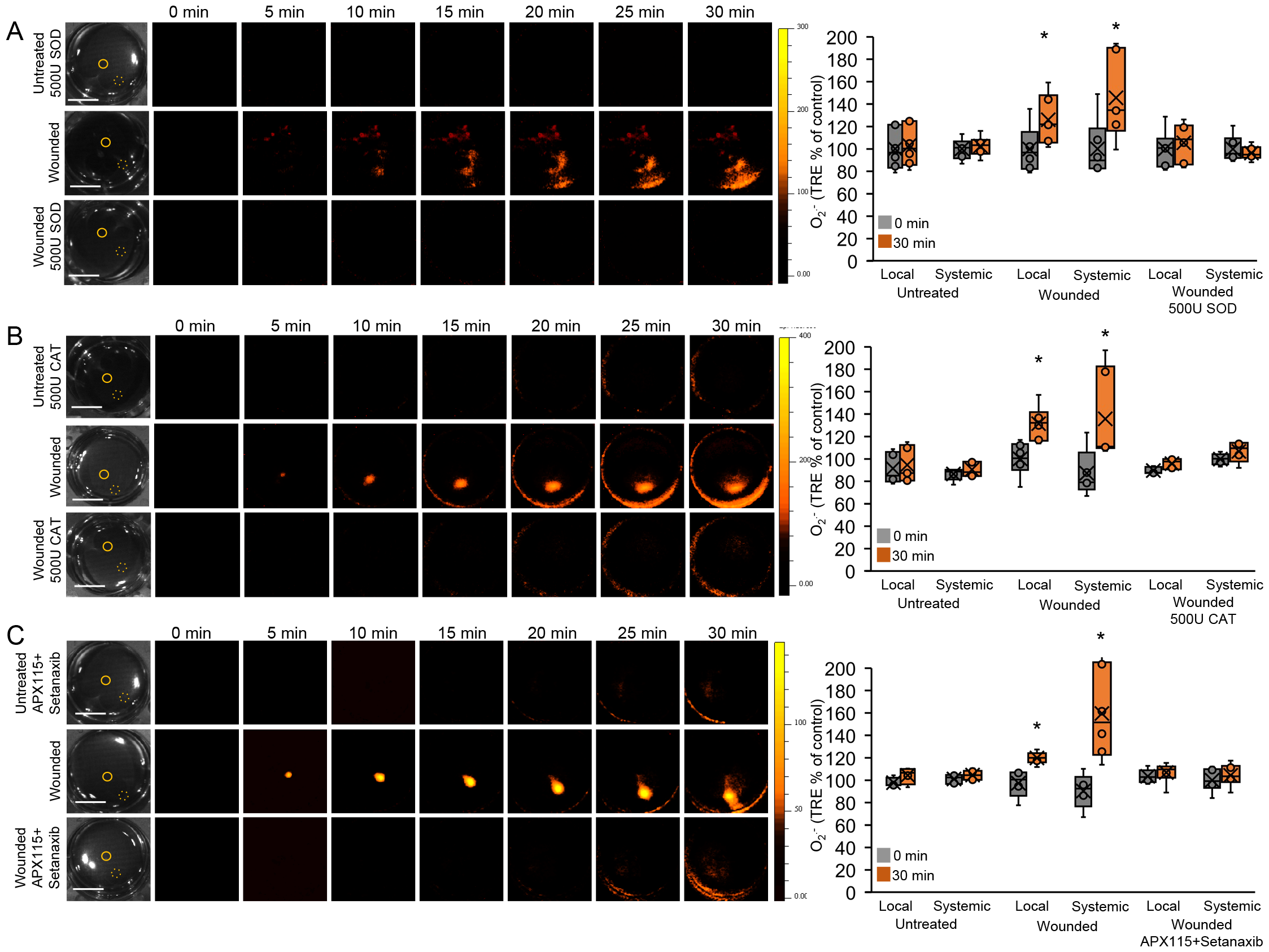
Inhibition of the wound induced O_2_·^-^ accumulation signal with catalase (CAT), superoxide dismutase (SOD), or a mixture of two NADPH oxidase (NOX) inhibitors. **A**. Monolayers of rat cardiomyocytes (H9c2) grown in culture were wounded with a heated metal rod to induce injury^20^ and O_2_·^-^ accumulation was imaged using DHE in the presence or absence of superoxide dismutase (added 60 min prior to wounding). Representative time-lapse images of whole plate O_2_·^-^ accumulation in treated and untreated plates are shown alongside bar graphs of combined data from all plates used for the analysis at the 0- and 30-min time points (local and systemic; Background levels of O_2_·^-^ measured at time 0, were subtracted from all other time points). Local and systemic sampling locations are indicated with solid and dashed circles, respectively. **B**. Same as in A, but instead of SOD, CAT was applied^20^. **C**. Same as in A, but instead of SOD, a mixture of two NOX inhibitors [APX115 (50 μM) and Setanaxib (50 μM)]^20^ was applied (application of APX115 or Setanaxib did not inhibit the O_2_·^-^ accumulation signal; Fig. S4). All experiments were repeated at least 3 times with 10 plates per experiment. Data is presented in A-C as box plot graphs; X is mean ± S.E., N=30, *P < 0.05, Student t-test. Scale bar, 1 cm. Abbreviations: CAT, catalase; DHE, dihydroethdium; NOX, NADPH oxidase; SOD, superoxide dismutase; TRE, total radiant efficiency.

### Enhanced cytosolic accumulation of H_2_O_2_ and changes in the redox sate of cells along the path of the H_2_O_2_ signal

We previously imaged the long-distance H_2_O_2_ signal in different organisms using various H_2_O_2_ /ROS-sensitive dyes^20^. These methods could however complicate the interpretation of results^2^. In addition, they did not measure changes in redox levels associated with the signal. We, therefore, generated monolayers of H9c2 rat cardiomyocyte cells expressing cytosolic roGFP2 (for redox), or cytosolic roGFP2-Orp1 (for H_2_O_2_ ), imaging (Figs. 2, S3). Analysis of wounded and unwounded H9c2 monolayers expressing these proteins revealed that the long-distance H_2_O_2_ signal is accompanied by enhanced cytosolic accumulation of H_2_O_2_ as well as changes in the redox state of cells along its path. These findings provide direct evidence for the putative function of the signal in altering the redox state of cells along its path, thereby, activating different redox responses (also supported by the identification of redox-response proteins in our proteomics analysis; Table S3).

### Production of extracellular superoxide and gap junction function are required for the propagation of the long-distance H_2_O_2_ signal

H_2_O_2_ -mediated cell-to-cell signaling in monolayers of H9c2 cells was previously shown to be blocked by catalase application, as well as a mixture of the NOX inhibitors APX115 and Setanaxib (but not each of these inhibitors applied individually)^20^. However, the role of superoxide in this signaling pathway was not substantiated. We, therefore, used external application of SOD, as well as imaging using dihydroethdium (DHE) in our experimental system. As shown in Figs. 1A and 3A, we could image the progression of the signal using DHE, and this signal was blocked by SOD or catalase application. In addition, a mixture of two NOX inhibitors blocked the signal imaged with DHE (Figs. 3B, S4). These findings are highly intriguing, as they support a model in which both superoxide and H_2_O_2_ are essential for the propagation of the long-distance H_2_O_2_ signal.

The ‘ROS wave’ long-distance H_2_O_2_ signal of plants was shown to require paracrine as well as juxtacrine signaling (the latter requiring the function of plasmodesmata, the plant equivalents of gap junctions)^18,20-26^. To test whether gap junction function is required for the long-distance H_2_O_2_ signal in H9c2 cells, we treated monolayers of H9c2 cells with CBX, βGA, or a mixture of the two^32^, prior to wounding. The combination of CBX+βGA, but not CBX or βGA blocked the progression of the H_2_O_2_ signal, imaged with DHE or PO1 (Figs. 4, S5). Our findings (Figs. 2, 4)^20^ support, therefore, a model in which paracrine and juxtacrine signaling, are both required for the propagation of the long-distance H_2_O_2_ pathway in mammalian cells (Fig. 4C). They further provide a potential mechanistic explanation to the directionality of the signal in H9c2 cells. While in other eukaryotic organisms, such as *Chlamydomonas* and MDA-MB-231 cells, the signal appears to diffuse away from the site of injury, without a clear direction, in H9c2 cells, it appears to be directional, following the orientation of cells in the monolayer^20^. This directionality could be related to the gap junctions that connect the cells and direct the signal along a specific path. In future studies, the polarity, and other aspects of the directionality of the signal, should be addressed, as these could play a key role in the function of the H_2_O_2_ signal in different tissues.

**Fig. 4.**
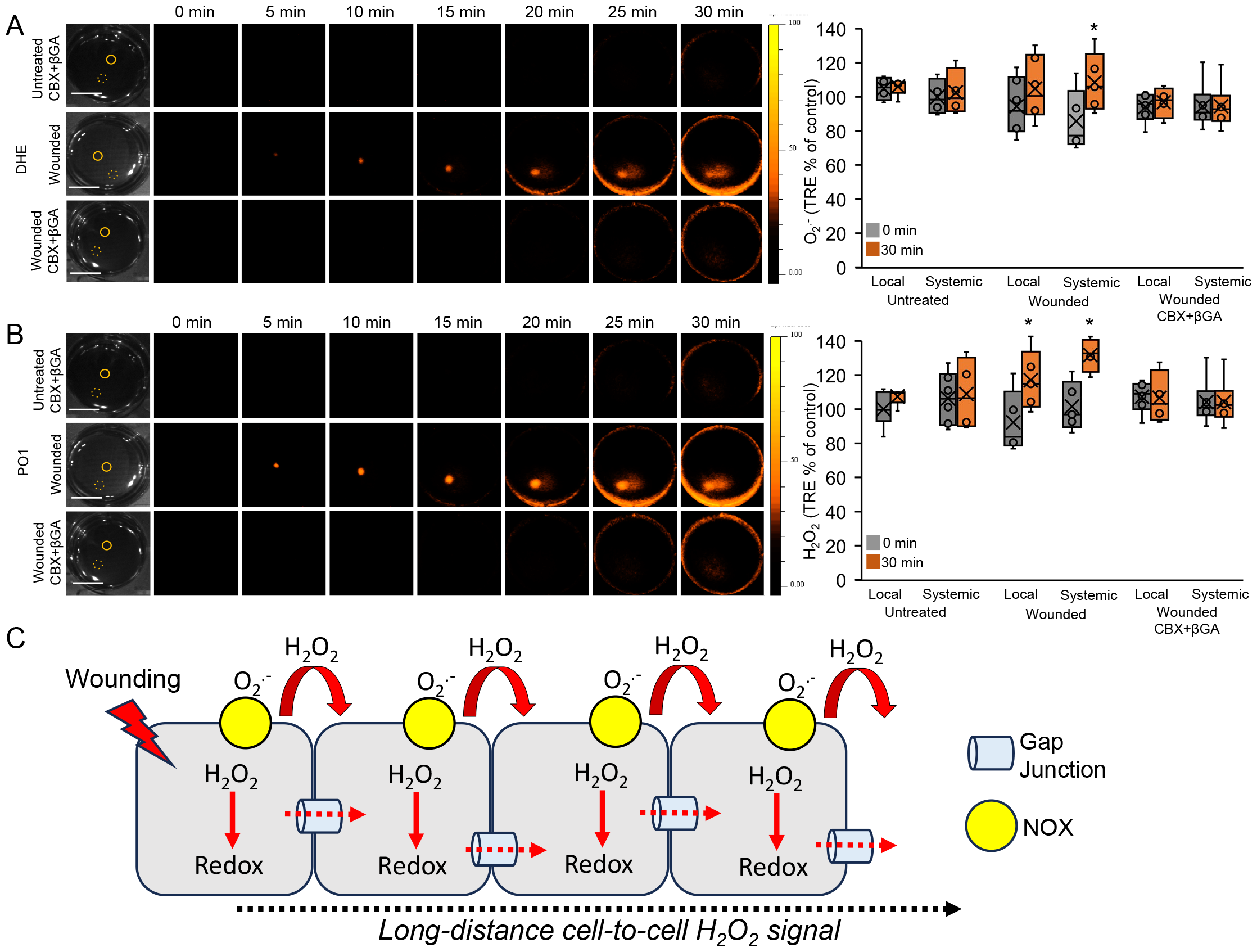
Inhibition of the wound induced H_2_O_2_ and O_2_·^-^ accumulation signal with gap junction inhibitors, and a model. **A**. Monolayers of rat cardiomyocytes (H9c2) grown in culture were wounded with a heated metal rod to induce injury^20^ and O_2_·^-^ accumulation was imaged using DHE in the presence or absence of a mixture of Carbenoxolone (CBX; 600 μM) and β-glycyrrhetinic acid (βGA; 180 μM), added 60 min prior to wounding. Representative time-lapse images of whole plate O_2_·^-^ accumulation in treated and untreated plates are shown alongside bar graphs of combined data from all plates used for the analysis at the 0- and 30-min time points (local and systemic; Background levels of O_2_·^-^ measured at time 0, were subtracted from all other time points). Local and systemic sampling locations are indicated with solid and dashed circles, respectively. **B**. Same as in A, but instead of DHE, PO1 (H_2_O_2_ accumulation) was used for imaging. **C**. A hypothetical model of the wound induced H_2_O_2_ and O_2_·^-^ accumulation cell-to-cell signal. Each sell along the path of the signal is shown to produce H_2_O_2_ and O_2_·^-^ propagating the signal across long distances in a process that requires NOX and gap junction function (*i*.*e*., a mixture of paracrine and juxtacrine signaling). See text for more details. All experiments were repeated at least 3 times with 10 plates per experiment. Data is presented in A and B as box plot graphs; X is mean ± S.E., N=30, *P < 0.05, Student t-test. Scale bar, 1 cm. Abbreviations: βGA, β-glycyrrhetinic acid; CBX, Carbenoxolone; DHE, dihydroethdium; NOX, NADPH oxidase; PO1, peroxy orange 1; TRE, total radiant efficiency.

## Summary and conclusions

Taken together, our findings reveal that the long-distance H_2_O_2_ cell-to-cell signaling pathway triggered by wounding in a monolayer of H9c2 cells (Fig. 4C) is accompanied by enhanced accumulation of H_2_O_2_ and altered redox state at the cytosol of cells along its path, as well as significant changes in the abundance of hundreds of proteins. In addition, we show that this pathway requires the production of superoxide as well as the function of gap junctions. As H_2_O_2_ signaling plays such a key role in mammalian physiology and development^1-3^, more studies are needed to dissect this pathway and determine how it is involved in different inflammatory responses, wound healing, development, cardiovascular disease, and other diseases/conditions.

## Conflict of interest

The authors declare no conflict of interest.

## Acknowledgments

pMF1707 was a gift from Dr. Marc Fransen (Addgene plasmid 125584) and pMOS016 was a gift from Dr. Adam Cohen (Addgene plasmid 163058).

## Funding

This work was supported by funding from the National Science Foundation (IOS-1932639, MCB-2224839), and the National Institute of Health grant (GM111364).

## Use of AI and AI-assisted technologies

None

## Author contribution

Y.F., L.R, and T.T.N. performed the experiments and/or analyzed the data. R.M., S.J.C., and Y.F. wrote the manuscript.

## Figure and Table Legends

**Table S1**. Proteins which expression is altered in local injured cells but not in systemic cells.

**Table S2**. Proteins which expression is altered in systemic cells but not in local injured cells.

**Table S3**. Proteins which expression is altered in both local injured cells and systemic cells.

**Fig. S1.**
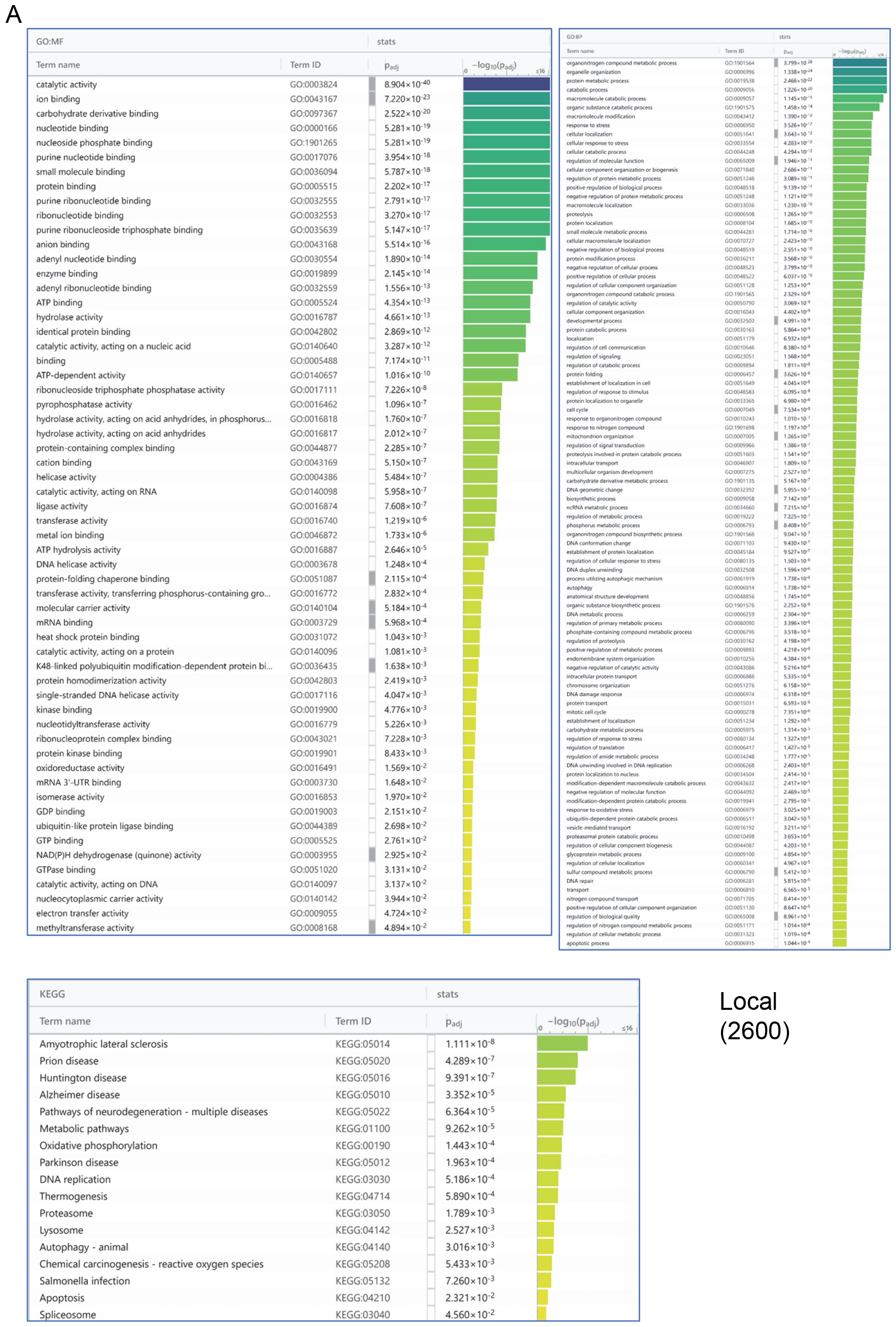
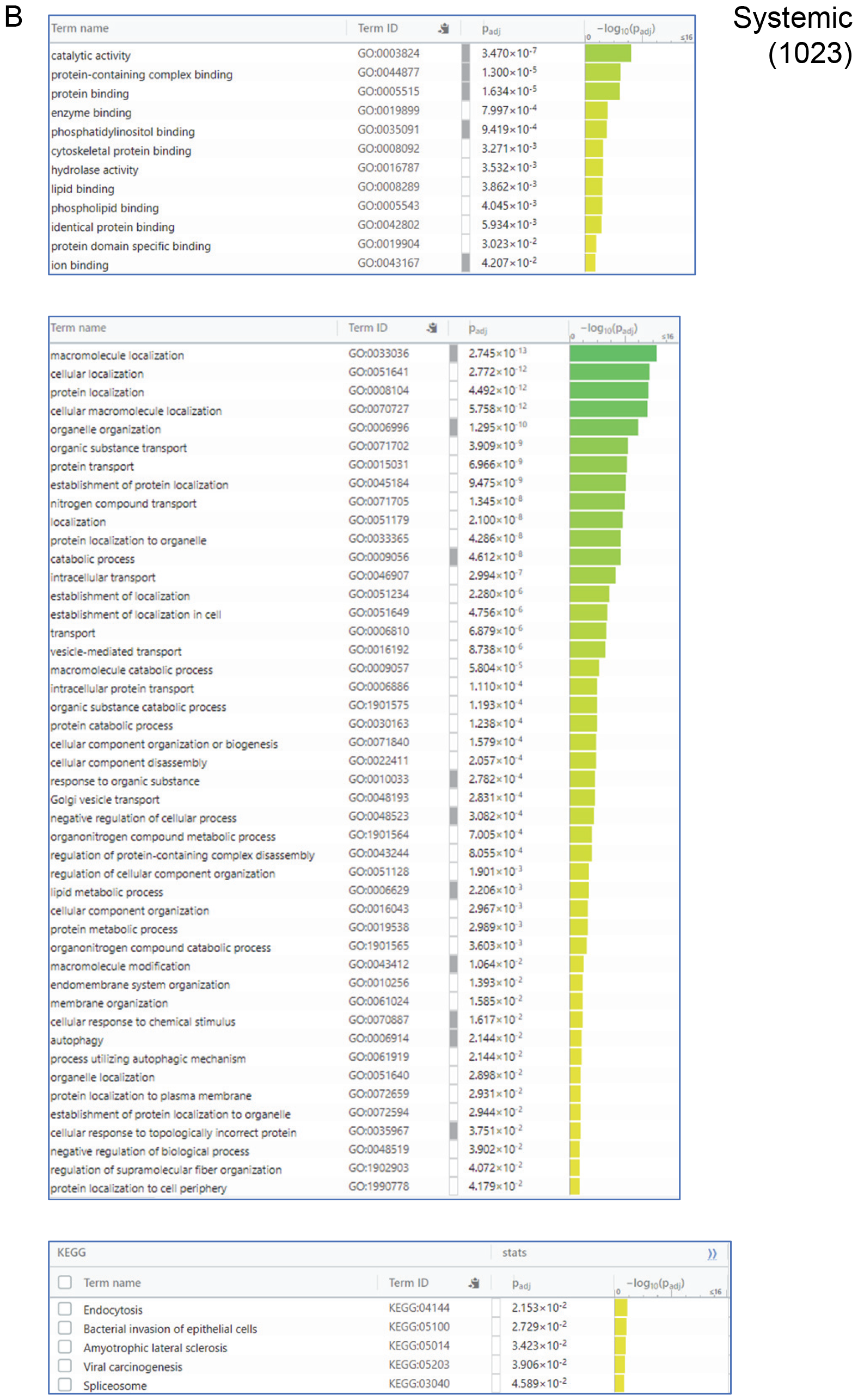
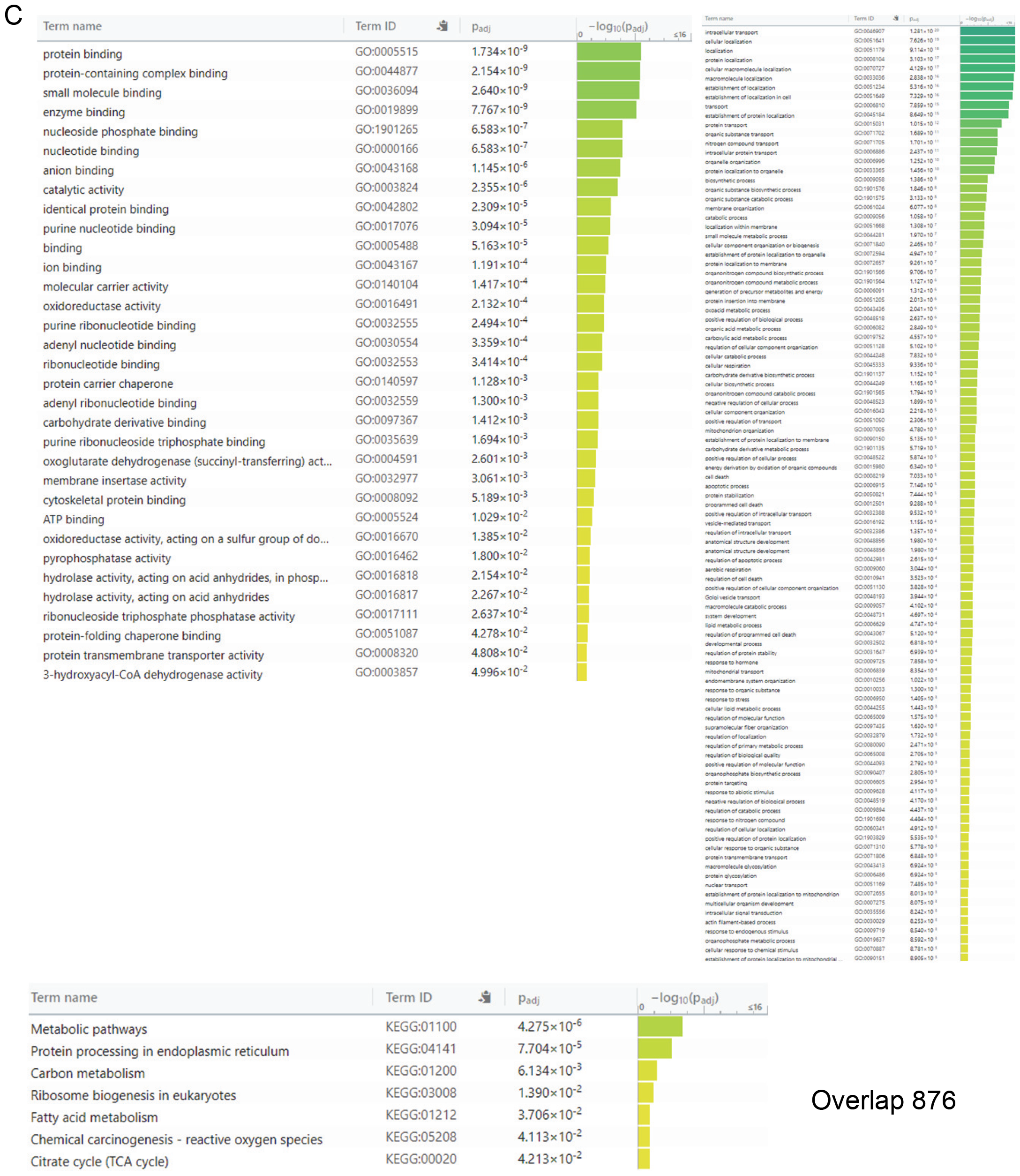
Gene Ontology (GO) and Kyoto Encyclopedia of Genes and Genomes (KEGG) annotation of the different proteins that accumulate in rat cardiomyocytes (H9c2) in local (**A**) and systemic (**B**) locations, as well as their overlap (**C**), in response to wounding. In support of Figs. 1A, 1B.

**Fig. S2.**
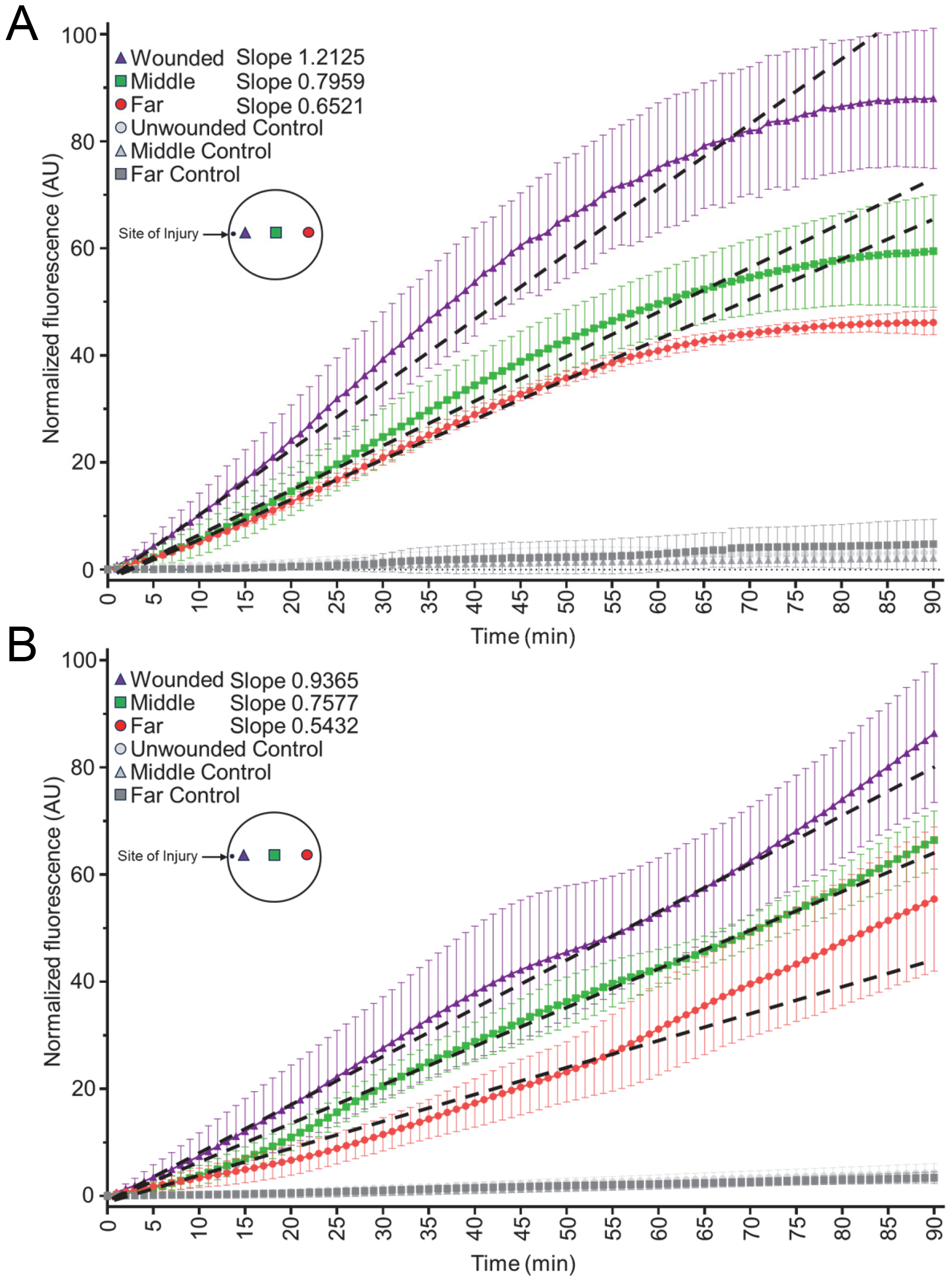
Accumulation of O_2_·^-^ and reactive oxygen species (ROS) imaged with dihydroethdium (DHE; **A**), or CellROX (**B**), at 3 different locations along the path of the H_2_O_2_ and O_2_·^-^ accumulation cell-to-cell signal. Rat cardiomyocytes (H9c2) grown in culture were wounded with a heated metal rod to induce injury^20^ and DHE or CellROX were used to image O_2_·^-^ and ROS accumulation at 3 different locations (wounded, middle, and far) in untreated or wounded cells using an automated Lionheart FX (BioTek) fluorescence microscope. Inserts in A and B show the location of the imaged cells within the well (96 well plate). All experiments were repeated at least 3 times with 10 wells per experiment. In support of Fig. 1A.

**Fig. S3.**
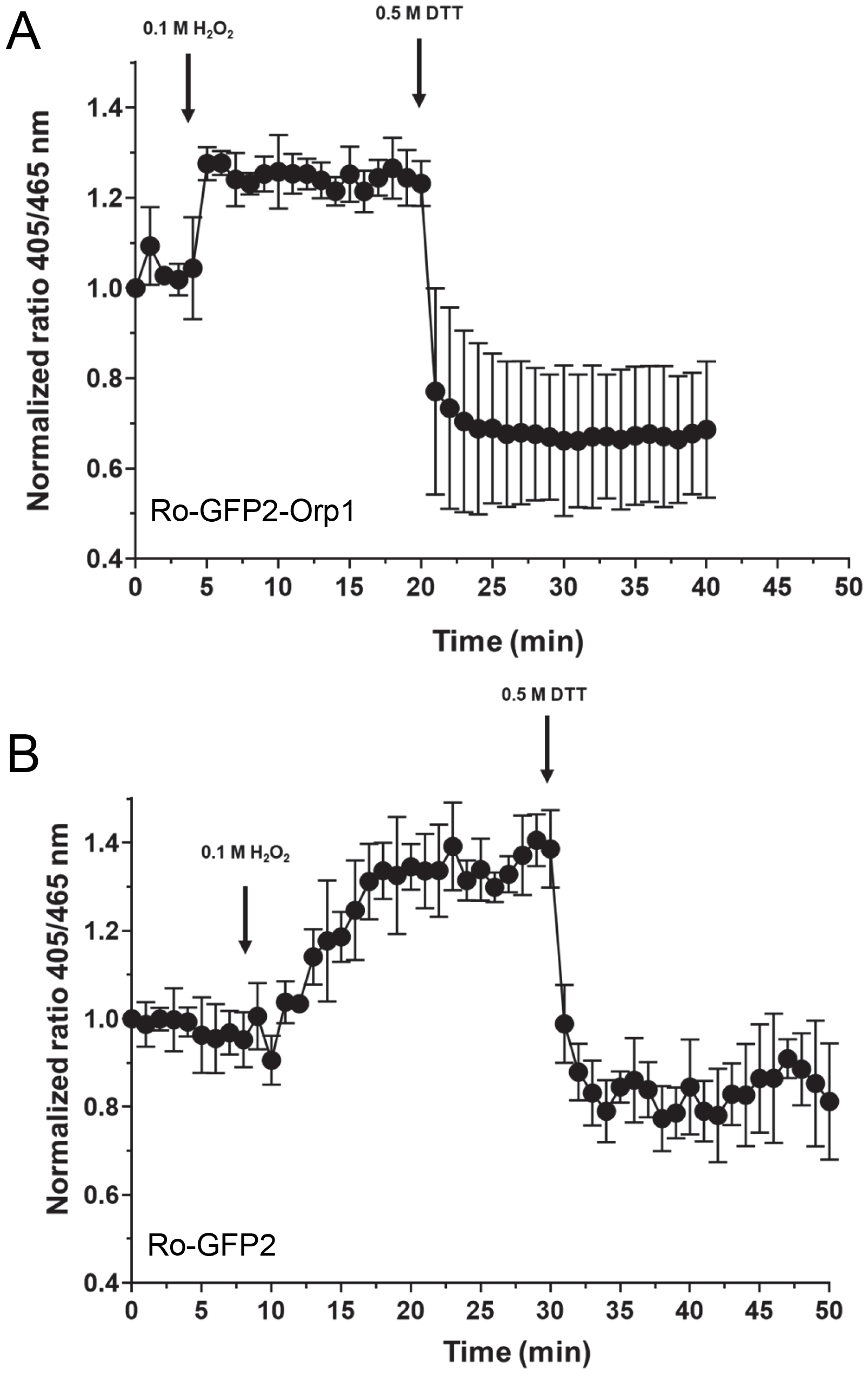
Calibration measurements of rat cardiomyocytes (H9c2) expressing roGFP2-Orp1 (**A**) or roGFP2 (**B**) and treated with H_2_O_2_ followed by DTT. After^24^. Cells were imaged using an automated Lionheart FX (BioTek) fluorescence microscope. In support of Fig. 2.

**Fig. S4.**
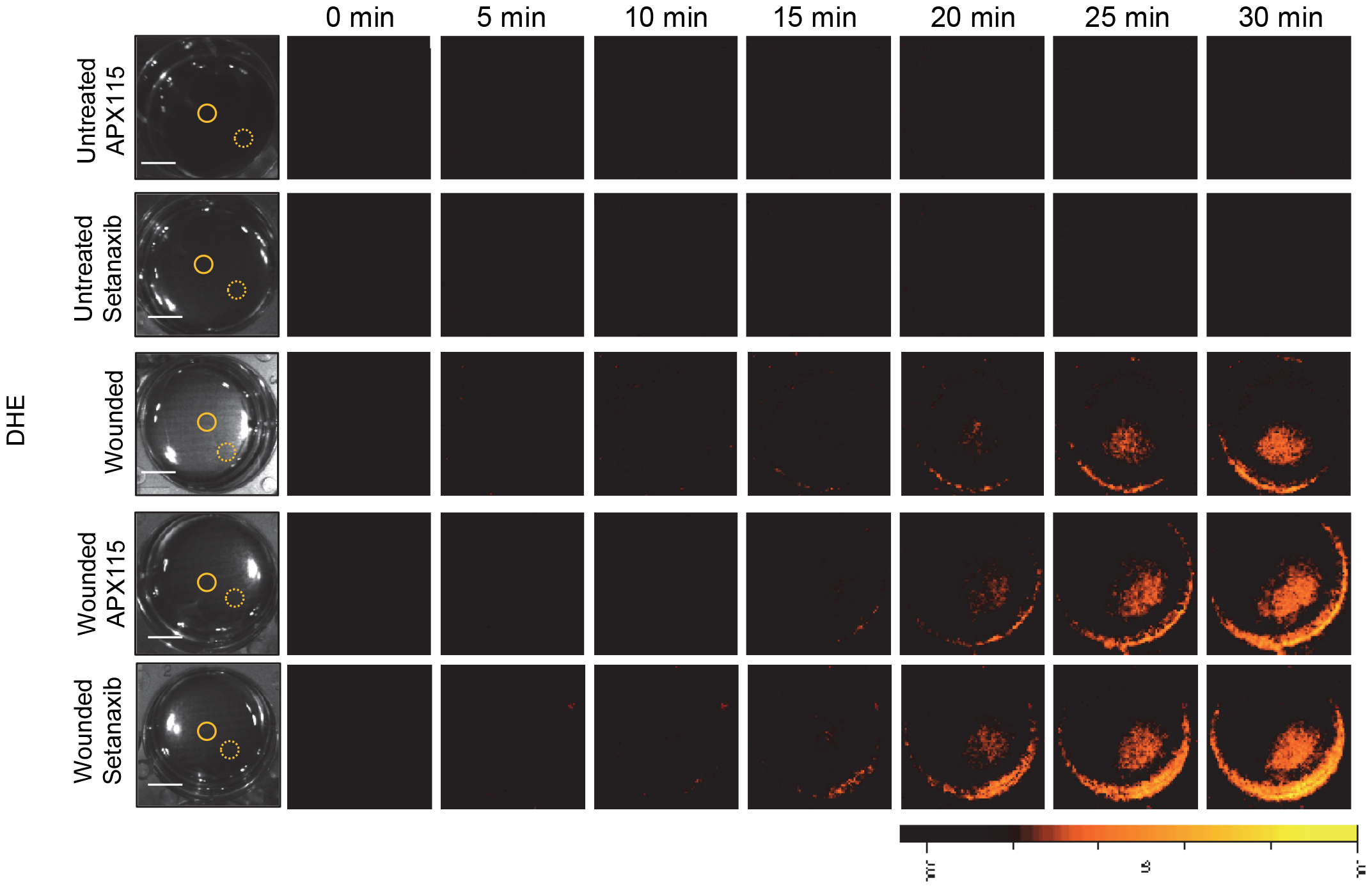
Application of APX115 (50 μM) or Setanaxib (50 μM) does not inhibit of the wound induced O_2_·^-^ accumulation signal. Monolayers of rat cardiomyocytes (H9c2) grown in culture were wounded with a heated metal rod to induce injury^20^ and O_2_·^-^ accumulation was imaged using DHE in the presence or absence of APX115 (50 μM) or Setanaxib (50 μM). Representative time-lapse images of whole plate O_2_·^-^ accumulation in treated and untreated plates are shown. Abbreviations: DHE, dihydroethdium. Scale bar, 1 cm. In support of Fig. 3.

**Fig. S5.**
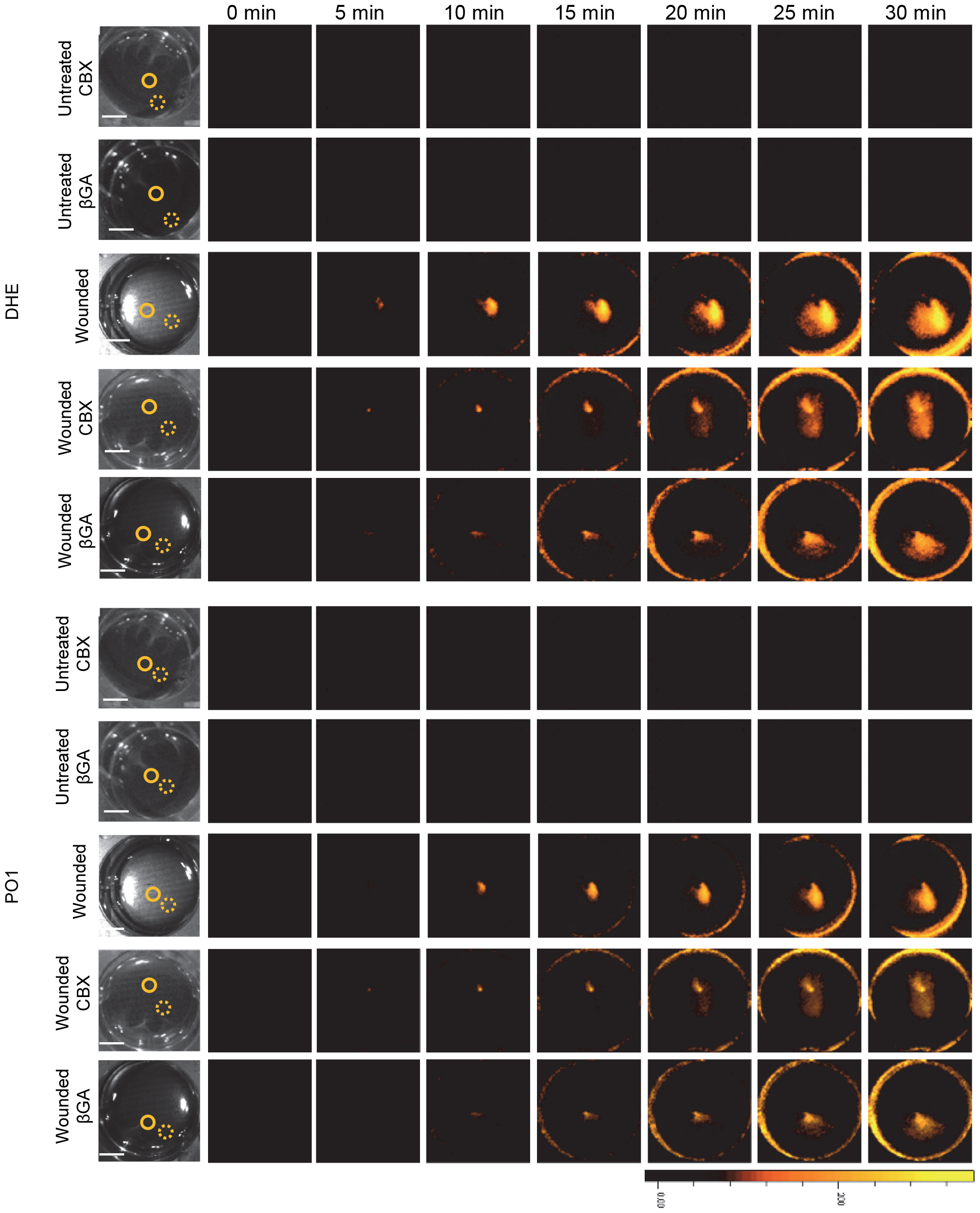
Application of Carbenoxolone (CBX) or β-glycyrrhetinic acid (βGA) does not inhibit of the wound induced H_2_O_2_ and O_2_·^-^ accumulation signal. Monolayers of rat cardiomyocytes (H9c2) grown in culture were wounded with a heated metal rod to induce injury^20^ and H_2_O_2_ and O_2_·^-^ accumulation were imaged with DHE (**A**) or PO1 (**B**) in the presence or absence of CBX (600 μM) or βGA (180 μM). Representative time-lapse images of whole plate O_2_·^-^ accumulation in treated and untreated plates are shown. Scale bar, 1 cm. Abbreviations: DHE, dihydroethdium; PO1, peroxy orange 1. In support of Fig. 4A, 4B.

## Notes

### Competing Interest Statement

The authors have declared no competing interest.

## References

1. Sies H, Jones DP. Reactive oxygen species (ROS) as pleiotropic physiological signalling agents. Nat Rev Mol Cell Biol. 2020 Jul;21(7):363–383. doi: 10.1038/s41580-020-0230-3.

2. Sies H, Belousov VV, Chandel NS, Davies MJ, Jones DP, Mann GE, Murphy MP, Yamamoto M, Winterbourn C. Defining roles of specific reactive oxygen species (ROS) in cell biology and physiology. Nat Rev Mol Cell Biol. 2022 Jul;23(7):499–515. doi: 10.1038/s41580-022-00456-z.

3. Sies H. Hydrogen peroxide as a central redox signaling molecule in physiological oxidative stress: Oxidative eustress. Redox Biol. 2017 Apr;11:613–619. doi: 10.1016/j.redox.2016.12.035.

4. Mittler R, Zandalinas SI, Fichman Y, Van Breusegem F. Reactive oxygen species signalling in plant stress responses. Nat Rev Mol Cell Biol. 2022 Oct;23(10):663–679. doi: 10.1038/s41580-022-00499-2.

5. Pató A, Bölcskei K, Donkó Á, Kaszás D, Boros M, Bodrogi L, Várady G, Pape VFS, Roux BT, Enyedi B, Helyes Z, Watt FM, Sirokmány G, Geiszt M. Hydrogen peroxide production by epidermal dual oxidase 1 regulates nociceptive sensory signals. Redox Biol. 2023 Jun;62:102670. doi: 10.1016/j.redox.2023.102670.

6. Enyedi B, Zana M, Donkó Á, Geiszt M. Spatial and temporal analysis of NADPH oxidase-generated hydrogen peroxide signals by novel fluorescent reporter proteins. Antioxid Redox Signal. 2013 Aug 20;19(6):523–34. doi: 10.1089/ars.2012.4594.

7. Grubisha MJ, Cifuentes ME, Hammes SR, Defranco DB. A local paracrine and endocrine network involving TGFβ, Cox-2, ROS, and estrogen receptor β influences reactive stromal cell regulation of prostate cancer cell motility. Mol Endocrinol. 2012 Jun;26(6):940–54. doi: 10.1210/me.2011-1371.

8. Xu M, Zhang Y, Xia M, Li XX, Ritter JK, Zhang F, Li PL. NAD(P)H oxidase-dependent intracellular and extracellular O2•-production in coronary arterial myocytes from CD38 knockout mice. Free Radic Biol Med. 2012 Jan 15;52(2):357–65. doi: 10.1016/j.freeradbiomed.2011.10.485.

9. Csányi G, Taylor WR, Pagano PJ. NOX and inflammation in the vascular adventitia. Free Radic Biol Med. 2009 Nov 1;47(9):1254–66. doi: 10.1016/j.freeradbiomed.2009.07.022.

10. Cadiz Diaz A, Schmidt NA, Yamazaki M, Hsieh CJ, Lisse TS, Rieger S. Coordinated NADPH oxidase/hydrogen peroxide functions regulate cutaneous sensory axon de- and regeneration. Proc Natl Acad Sci U S A. 2022 Jul 26;119(30):e2115009119. doi: 10.1073/pnas.2115009119.

11. Evans IR, Rodrigues FS, Armitage EL, Wood W. Draper/CED-1 mediates an ancient damage response to control inflammatory blood cell migration in vivo. Curr Biol. 2015 Jun 15;25(12):1606–12. doi: 10.1016/j.cub.2015.04.037.

12. Petersen SV, Poulsen NB, Linneberg Matthiesen C, Vilhardt F. Novel and Converging Ways of NOX2 and SOD3 in Trafficking and Redox Signaling in Macrophages. Antioxidants (Basel). 2021 Jan 25;10(2):172. doi: 10.3390/antiox10020172. Vesicles

13. Amblard I, Thauvin M, Rampon C, Queguiner I, Pak VV, Belousov V, Prochiantz A, Volovitch M, Joliot A, Vriz S. H_2_ O_2_ and Engrailed 2 paracrine activity synergize to shape the zebrafish optic tectum. Commun Biol. 2020 Sep 29;3(1):536. doi: 10.1038/s42003-020-01268-7.

14. Buck T, Hack CT, Berg D, Berg U, Kunz L, Mayerhofer A. The NADPH oxidase 4 is a major source of hydrogen peroxide in human granulosa-lutein and granulosa tumor cells. Sci Rep. 2019 Mar 5;9(1):3585. doi: 10.1038/s41598-019-40329-8.

15. Niethammer P, Grabher C, Look AT, Mitchison TJ. A tissue-scale gradient of hydrogen peroxide mediates rapid wound detection in zebrafish. Nature. 2009 Jun 18;459(7249):996–9. doi: 10.1038/nature08119.

16. Razzell W, Evans IR, Martin P, Wood W. Calcium flashes orchestrate the wound inflammatory response through DUOX activation and hydrogen peroxide release. Curr Biol. 2013 Mar 4;23(5):424–9. doi: 10.1016/j.cub.2013.01.058.

17. Katikaneni A, Jelcic M, Gerlach GF, Ma Y, Overholtzer M, Niethammer P. Lipid peroxidation regulates long-range wound detection through 5-lipoxygenase in zebrafish. Nat Cell Biol. 2020 Sep;22(9):1049–1055. doi: 10.1038/s41556-020-0564-2.

18. Fichman Y, Zandalinas SI, Peck S, Luan S, Mittler R. HPCA1 is required for systemic reactive oxygen species and calcium cell-to-cell signaling and plant acclimation to stress. Plant Cell. 2022 Oct 27;34(11):4453–4471. doi: 10.1093/plcell/koac241.

19. Sousa T, Gouveia M, Travasso RDM, Salvador A. How abundant are superoxide and hydrogen peroxide in the vasculature lumen, how far can they reach? Redox Biol. 2022 Dec;58:102527. doi: 10.1016/j.redox.2022.102527.

20. Fichman Y, Rowland L, Oliver MJ, Mittler R. ROS are evolutionary conserved cell-to-cell stress signals. Proc Natl Acad Sci U S A. 2023 Aug;120(31):e2305496120. doi: 10.1073/pnas.2305496120.

21. Peláez-Vico MÁ, Fichman Y, Zandalinas SI, Van Breusegem F, Karpiński SM, Mittler R. ROS and redox regulation of cell-to-cell and systemic signaling in plants during stress. Free Radic Biol Med. 2022 Nov 20;193(Pt 1):354–362. doi: 10.1016/j.freeradbiomed.2022.10.305.

22. Zandalinas SI, Fichman Y, Mittler R. Vascular Bundles Mediate Systemic Reactive Oxygen Signaling during Light Stress. Plant Cell. 2020 Nov;32(11):3425–3435. doi: 10.1105/tpc.20.00453.

23. Fichman Y, Myers RJ Jr, Grant DG, Mittler R. Plasmodesmata-localized proteins and ROS orchestrate light-induced rapid systemic signaling in Arabidopsis. Sci Signal. 2021 Feb 23;14(671):eabf0322. doi: 10.1126/scisignal.abf0322.

24. Fichman Y, Mittler R. A systemic whole-plant change in redox levels accompanies the rapid systemic response to wounding. Plant Physiol. 2021 May 27;186(1):4–8. doi: 10.1093/plphys/kiab022.

25. Miller G, Schlauch K, Tam R, Cortes D, Torres MA, Shulaev V, Dangl JL, Mittler R. The plant NADPH oxidase RBOHD mediates rapid systemic signaling in response to diverse stimuli. Sci Signal. 2009 Aug 18;2(84):ra45. doi: 10.1126/scisignal.2000448.

26. Zandalinas SI, Fichman Y, Devireddy AR, Sengupta S, Azad RK, Mittler R. Systemic signaling during abiotic stress combination in plants. Proc Natl Acad Sci U S A. 2020 Jun 16;117(24):13810–13820. doi: 10.1073/pnas.2005077117.

27. Rowland L, Marjault HB, Karmi O, Grant D, Webb LJ, Friedler A, Nechushtai R, Elber R, Mittler R. A combination of a cell penetrating peptide and a protein translation inhibitor kills metastatic breast cancer cells. Cell Death Discov. 2023 Aug 31;9(1):325. doi: 10.1038/s41420-023-01627-3.

28. Karmi O, Sohn YS, Zandalinas SI, Rowland L, King SD, Nechushtai R, Mittler R. Disrupting CISD2 function in cancer cells primarily impacts mitochondrial labile iron levels and triggers TXNIP expression. Free Radic Biol Med. 2021 Nov 20;176:92–104. doi: 10.1016/j.freeradbiomed.2021.09.013. Transfection, proteomics

29. Karmi O, Rowland L, King SD, Manrique-Acevedo C, Cabantchik IZ, Nechushtai R, Mittler R. The [2Fe-2S] protein CISD2 plays a key role in preventing iron accumulation in cardiomyocytes. FEBS Lett. 2022 Mar;596(6):747–761. doi: 10.1002/1873-3468.14277. Proteomics

30. Sohn YS, Tamir S, Song L, Michaeli D, Matouk I, Conlan AR, Harir Y, Holt SH, Shulaev V, Paddock ML, Hochberg A, Cabanchick IZ, Onuchic JN, Jennings PA, Nechushtai R, Mittler R. NAF-1 and mitoNEET are central to human breast cancer proliferation by maintaining mitochondrial homeostasis and promoting tumor growth. Proc Natl Acad Sci U S A. 2013 Sep 3;110(36):14676–81. doi: 10.1073/pnas.1313198110. DHE imaging

31. Lisse TS, King BL, Rieger S. Comparative transcriptomic profiling of hydrogen peroxide signaling networks in zebrafish and human keratinocytes: Implications toward conservation, migration and wound healing. Sci Rep. 2016 Feb 5;6:20328. doi: 10.1038/srep20328.

32. Behringer EJ, Socha MJ, Polo-Parada L, Segal SS. Electrical conduction along endothelial cell tubes from mouse feed arteries: confounding actions of glycyrrhetinic acid derivatives. Br J Pharmacol. 2012 May;166(2):774–87. doi: 10.1111/j.1476-5381.2011.01814.x.

